# Factors affecting secondary sex characteristics in the yellowtail tetra *Astyanax altiparanae*

**DOI:** 10.1101/698100

**Authors:** Diógenes H. Siqueira-Silva, Rafaela M. Bertolini, Nycolas L. Pereira, Nivaldo F. Nascimento, José A. Senhorini, Lucas Henrique Piva, José Bento S. Ferraz, George S. Yasui

**Author notes:** Correspondence: George Shigueki Yasui.

## Abstract

This work aimed to analyze factors affecting secondary sexual characteristics in the yellowtail tetra *Astyanax altiparanae.* For this, seventy-five specimens were first separated into three different size classes (small, medium and large groups) between two seasons, summer and winter. In all groups, male fish were consistently bigger in the summer. On the other hand, females from both seasons presented in media the same length into the length classes. Afterwards, we performed histological analyses of the gonads to first confirm the genus and investigate the phase of maturation of each animal. During the winter, most of the small animals were males (22), most of the large animals, females (23), and the medium size animals followed a tendency of 1:1 ratio (9 male: 16 female). In the summer, male were the majority in both small (20) and medium (20) size. Larger-size animals were female (23). Then, in order to analyze the influence of genus, phase of maturation, season of the year, the number, and length of the animals spinelets, we diaphanized, counted, and measured them in each animal. Our results demonstrated that the spinelets are a sexual secondary characteristic of male genus independently of the size, season and phase of maturation. However, some tendencies were observed. Males bigger than 48 mm always presented spinelets; their size are in media the double in summer in comparison to winter; and summer males presents more rays with spinelets in the summer. Curiously, the larger specimen sampled was a female presenting spinelets in five rays. Lastly, we performed the gonadectomization of the animals and hypothesized that gonad hormones will directly influence this characteristic. The gonadectomization only initially influence on the size and number of spinelets in the anal fin rays, since the thirty-day-gonadectomized animals presented few and smaller spinelets against the control ones. However, the spinelets normalized in ninety-day-gonadectomized specimens. Such a work showed spinelets can be considered a secondary sexual characteristic to distinct male from female and can be used in the management in specimens bigger than 48 mm, but cannot indicate fish sterility.

**Summary statement:** This study elucidated whether the size, sex, environmental conditions, and gonadal development affect the development of spinelets, a bony structure presented in anal fins in mature fish. Additionally, gonadectomized fish were used to elucidate the effect of gonad on the rise of such structures. Interesting new data showed that such a secondary sex characteristic is influenced by sex, size, gonadal development, and season of the year, but spinelets arose even within gonadectomized fish; this suggests that such a structure is not indicative of sterility in this species.

## Introduction

Most teleost fish spawning is a seasonal event that depends on a series of environmental conditions that triggers reproductive migration and subsequent gonadal maturation. Since photoperiod, rainfall, and temperature improve food availability for the offspring, maturation occurs when there is more rainfall and humidity present (Baldisserotto, 2013). On the other hand, species such the yellowtail tetra *Astyanax altiparanae* presents intertidal spawning and may reproduce several times along the year (Orsi et al., 2004), including the winter season.

The objective of sexual reproduction is to proceed gametogenesis, which leads to fertilization; however, changes in behavior or the hypertrophy of specific structures are concomitant to these processes. They may also influence the success rate of fish reproduction, reflecting on both their physiology and behavior. For example, female red snapper *Lutjanus campechanus* are vastly more productive than smaller specimens (Bohnsack, 1998) and, in the black rockfish *Sebastes melanops*, older female produce larvae with a higher growth rate and more resistant to starving (Berkeley et al., 2004).

In some species of teleost, specific secondary sexual characteristics, which are the result from sexual dimorphism, can influence the reproduction and suggest the beginning of the reproductive period. In the Amazonian peacock bass *Cichla sp.*, males develop a post occipital protuberance during the reproductive period (Kelber, 1999). Such a structure is composed basically of lipids, which is supposed to provide energy during parental care (Jepsen et al., 1999; Muñoz et al., 2006). In several species from Characiforms order, it is known that the mature male develops spinelets in the anal fin during the reproductive period in order to adhere to female anal fin during spawning (Malabarba and Weitzman, 2003). Such structures may be applied in aquaculture practice since it facilitates sexual reproduction prior to spawning procedures. However, although the development of secondary sex characteristics is often associated with gonadal maturation, there is no confirmation that the rise of spinelets are indeed predictors of gonadal maturation.

Thus, the objective of the study was to understand the physiological role of the spinelets to *A. altiparanae* reproduction as well as to help fish broodstock management. The present study evaluated: the growth dynamic of spinelets in accordance to the genre and length of the animals during Winter and Summer seasons and gonadectomization effects on the spinelets’ dynamic. The choice to use this species for such a study was motivated by its good livestock characteristics, such as ease of handling, small size, early sexual maturity, and intertidal spawning (Garuti, 2003).

## Materials and Methods

### Origin of broodstock and gamete sampling

All the procedures were performed in line with the Ethical Committee for the Care and Use of Laboratory Animals in Chico Mendes Institute (CEUA - CEPTA #02031.000033 / 2015-11).

The yellowtail tetra *A. altiparanae* used in this study were collected from Mogi Guassu River (21.925706 S, 47.369496 W) and maintained in 1000 m^2^ earthen ponds (≈ 500 fish per tank, SL ≈ 12 cm). As this species spawns spontaneously, F1 offspring was produced within few months, and those fish were used in the experiments.

### Experiment I – Secondary sex characteristics in winter and summer seasons

This Experiment was divided into two trials. The first one was performed during the winter (July 2013) and the second one during the summer (January 2014), each resulting in a reproductive peak and rest period. In each period, 75 specimens were collected and divided into three groups of 25 fish each according to their total length. For this, the fish were euthanized in menthol solution (100 mg.L^-1^) and had the standard length (mm), height (mm), and weight (g) registered, being categorized as small-sized-group (45.85 ± 5.39 mm), medium-sized-group (60.54 ± 5.89 mm), and large-sized-group (80.72 ± 7.08 mm) (Fig. 1).

**Figure 1.**
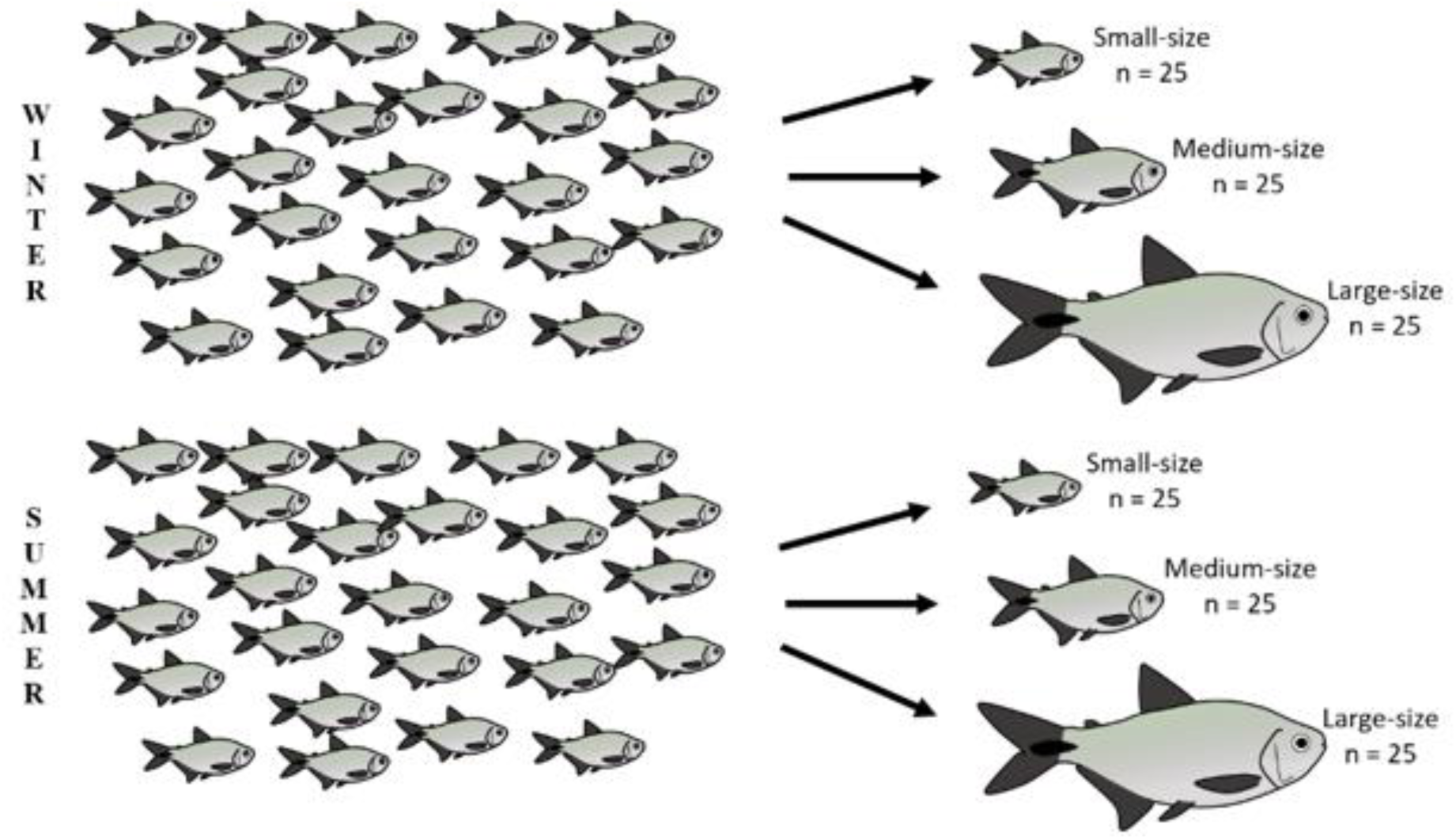
Experiment I. Secondary sexual morphological characterization of the animals in the winter and summer seasons

#### Histological analysis of the gonads

To define the sex and maturation status of the collected specimens, fish were anesthetized using 2-Phenoxy-Ethanol (C_6_H_10_O_2_ – SIGMA-ALDRICH), their gonads were removed, cut into transverse and longitudinal sections, and fixed with Bouin’s fixative (Adria laboratories, Londrina, Brazil) for 24 hours. Samples were dehydrated in a series of graded ethanol, embedded in paraffin – polyisobutylene mixture (Paraplast®, SIGMA-ALDRICH), sectioned at 5.0 μm on a microtome (Leica RM2235, Wetzlar, Germany) equipped with steel blade (Leica 818), and the sections were then stained with hematoxylin and eosin. All material was examined on a microscope (Nikon-Eclipse CI, Japan); digital images were captured with a CCD camera (Nikon DSF1, Nikon, Tokyo, Japan) and analyzed with NIS-Elements AR software (Nikon, Tokyo, Japan). Gonadal maturation was classified according with the criteria adopted by Brown-Peterson et al. (2011).

#### Morphological analysis of the diaphanized fins rays

In order to determine the number and length of fin rays and the presence, number, and length of the spinelets, the anal fin of each animal was removed and fixed in 5% formalin for 24 h, being posteriorly submitted to diaphanization using the Potthoff (1984) protocol. The samples were washed in distilled water for 24 hours and then dehydrated in ethanol solutions (50% for 24 hours and 95% for 24 hours). For the cartilage staining, the samples were incubated in Alcian blue acidified ethanol solution (20 mg of Alcian blue 200 mg.L^-1^, 60% of ethanol, 40% of glacial acetic acid) for 24 hours. After this, the samples were incubated in saturated borate solution (5 hours), and subsequently, in whitening solution (3% of H_2_O_2_ and 2% of KOH, 2 hours), and clarifying solution (35% saturated borate solution, and 65% of the whitening solution, 7 hours). For bone staining, the samples were then incubated in Alizarin solution (Alizarin 2% and KOH 2% in distilled water, for 24 hours) and transferred to successive preservation solutions (solution I: glycerin 30% and 70% of KOH 2% solution for 24 hours; solution II: glycerin 60% and 40% of KOH 2% solution). Lastly, the samples were kept in maintenance solution (glycerin added with thymol).

The diaphanized fins were examined under a stereomicroscope (Nikon SMZ 1500, Nikon, Tokyo, Japan), and digital images were captured using a CCD camera (Nikon DSF1, Nikon, Tokyo, Japan). The images were analyzed with NIS-Elements AR software (Nikon, Tokyo, Japan). The whole fish images were performed using a professional camera (Nikon 3100, Tokyo, Japan).

### Experiment II – Effects of the gonadectomy on the secondary sex characteristics

#### Long-term anesthesia for surgical procedures

Prior to surgery, fish were placed in 2 L-beakers containing 1 Liter of 0.7% of 2-phenoxyethanol solution in tap water. They had the following anesthetics parameters assessed: hyperactivity, erratic swimming, non-physiologic positioning, and absence of opercular ventilation.

After the anesthesia induction, fish were transferred to a surgical apparatus as described by Harms and Lewbart (2000). Briefly, it was composed by a 6 L-aquaria containing anesthetic solution and a pump (4 mL.s^-1^) coupled to a silicon cannula (3.5 mm internal diameters and 4.5 mm external diameter). The fish were placed in dorsal decubitus on a sponge cut in “V tray” supported by a net placed in the water surface. The cannula was inserted into the oral cavity of the fish in a way that kept the solution passing by the gills and exiting through the operculum.

Twenty-four fish were divided into six treatments (n = 4): for T1, the control treatment, a solution free of any anesthetic was used; for T2, T3, T4, T5, and T6, solutions containing 0.1, 0.2, 0.3, 0.4, and 0.5% of 2-phenoxyethanol were used. For each treatment, the period of anesthesia induction was registered, from the ending of operculum movement until the return of the caudal reflexes. As a reference, 60 minutes was adopted as the maximum period of anesthesia, in which it is possible to perform surgical interventions of long durations (Yasui et al., 2009).

#### Gonadectomy

Before the surgery, the fish were fasted for 24 hours in order to empty their gastrointestinal system. During the surgery, the fish were anesthetized and positioned in surgical beds as previously described. As determined by the previous experiment, a maintenance anesthetic solution containing 0.5% of 2-phenoxyethanol was used. A ventral incision was made from the pectoral fins to the posterior region of the pelvic fins. The coelomic cavity was explored in order to find the gonads, which were extracted with a hemostatic clamp, without the necessity of suture. The coelomic cavity was sutured with two separated single stiches using a class II-monophilamentar wire (nylon 4-0, Shalon fios cirúrgicos LTDA., Brazil), with approximated distance of 9 mm (Yasui et al., 2009).

Fourteen specimens presenting spicules, evident to touch, were separated in two groups (n = 7): in group 1 (control), the fish were just incised and then sutured; and in group 2, the gonadectomy was properly performed. To standardize the surgical duration in both treatments, the non-gonadectomized fish were left in the surgical bed during the same time as the gonadectomized.

After the surgery, the fish were allocated in a 416 L-aquarium, free from light (covered in aluminum foil), in water containing 2 g of Aurotrim (Chlortetracycline 1.5%, Sulfadiazine 7.5% and Trimethoprim 1.5%) for each 1000 L, deprived of feed for 96 hours. The stitches were removed after 4 days post-surgery (4 dps). After 30 dps, the fish were captured, killed by anesthesia, and submitted to diaphanization of the anal fin as previously described.

#### Statistics

All data are presented as mean ± standard deviation. The data were tested for normality using Liliefors, and in the case of normal distribution, were submitted to ANOVA followed by Tukey’s multiple range test. For the comparison of two means (e.g. winter and summer, gonadectomized or non-gonadectomized), a non-paired T-test was used. In all cases, the α = 5%.

## Results

### Experiment I

#### Histological analysis of the gonads

##### Winter Trial

Most of the specimens categorized as small size were male. The majority was in the developing stage (11 in the Initial Maturation; 9 in Mid Maturation) (Table 1; Fig. 2). Female were initiating the developing phases (Table 1; Fig. 3). One individual could not be identified. Most of medium-size fish were female. There were Immature, in Developing, and Mature animals (Table 1; Fig. 3). All the male specimens were in Developing stage; three of them were initiating the maturation and six in mid maturation (Table 1). From large size fish, 23 were female, in Developing phases and Mature, the majority (Table 1). Both of the male were in Developing (Table 1).

**Table I.** Reproductive Classes of the individuals sampled by size in year station.

| Animals |  | Winter |  |  | Summer |  |  |  |
| --- | --- | --- | --- | --- | --- | --- | --- | --- |
| size | Sex | Immature | Developing | Spawning Capable | Immature | Developing | Spawning Capable | Regressing |
| Small | Male | 1 | 20 | 1 | 1 | 16 | 2 | 1 |
|  | Female | 0 | 2 | 0 | 0 | 4 | 1 | 0 |
| Medium | Male | 0 | 9 | 0 | 0 | 15 | 4 | 1 |
|  | Female | 1 | 11 | 4 | 0 | 0 | 5 | 0 |
| Big | Male | 0 | 2 | 0 | 0 | 1 | 0 | 0 |
|  | Female | 0 | 9 | 14 | 4 | 3 | 16 | 0 |

**Figure 2.**
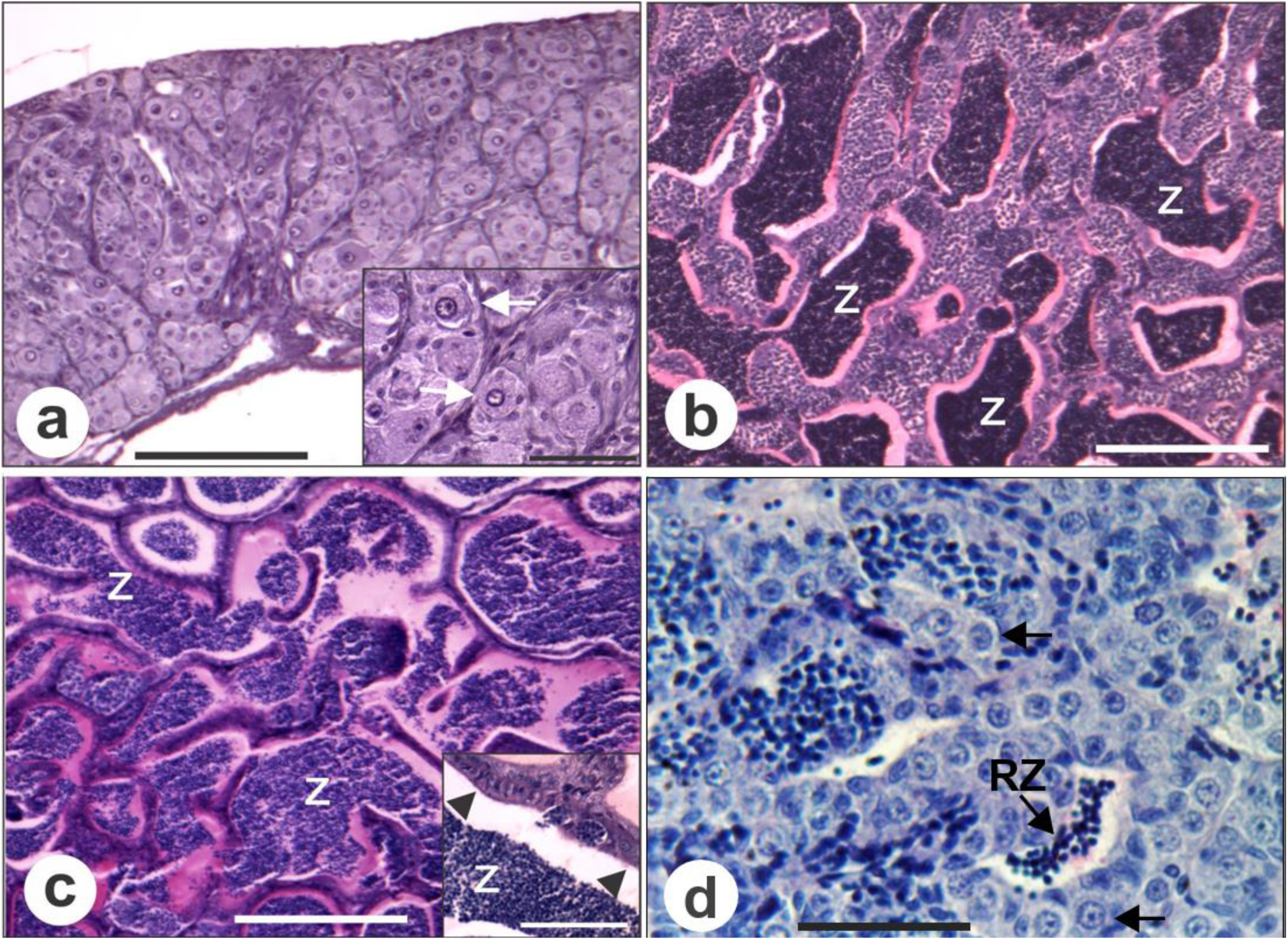
Histology characteristics of *A. altiparanae* testes phases. a) Immature; Inset: highlighting spermatogonia (arrow), which are very characteristic in this phase b) In Developing; c) Spawning capable; Inset: showing discontinuous epithelium (arrow head) d) Regressing. Spermatogonia (arrow), RZ = Residual spermatozoa. Scale bars: a-d 100 mm; Insets: 25 μm.

**Figure 3.**
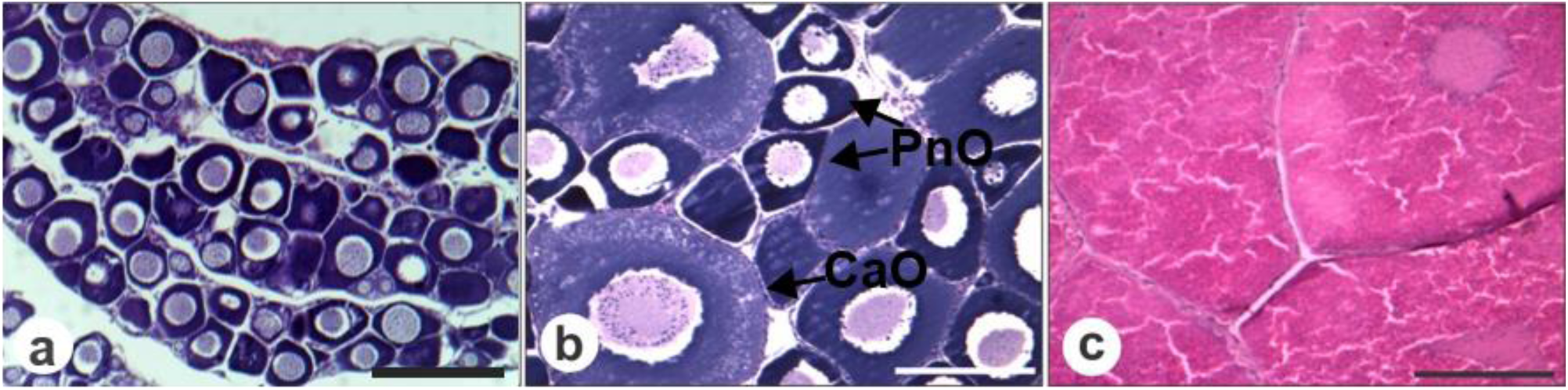
Histology characteristics of *A. altiparanae* ovaries phases. a) Immature, showing only Oocytes with perinucleolar nucleous; b) In Developing, showing pre-vitellogenic oocytes with perinuclear nucleolus (PnO) and oocytes with Cortical alveoli (CaO); c) Spawning capable; with vitellogenic oocytes. Scale bars: a-c 100 μm.

##### Summer Trial

In both of the small and medium size categories, most of the specimens were male (Table 1). Among small size fish, the majority were in Developing phases. Among male, five were in Initial Maturation and 11 in Mid Maturation. Most of the medium size males were also in Mid Maturation, and all the females were Mature (Table 1). Among specimens categorized as large size length, most were female. In that category, fish were most Mature (Table 1; Figs. 2-3).

##### Morphological analysis of the diaphanized fins rays

The male captured in winter presented a decreased number, length, and distribution of ray fin spinelets, with lower GSI when compared with fish captured in summer (Table 2). In winter, 23 out of 37 (62.1%) captured males presented spinelets while all 37 (100%) captured male presented spinelets in summer (Fig. 4).

**Table II.** Meristic and somatic parameters of *Astyanax altiparanae* male in the Winter 2013 and Summer 2014.

| Season | Size classes | Number of animals (n) | Weight (g) | Pattern length (mm) | GSI (%) | Spinelets presence (%) | Spinelets length (mm) | Total of spinelets | Rays with spinelets |
| --- | --- | --- | --- | --- | --- | --- | --- | --- | --- |
| Winter | Small | 22 | 2.79 ± 0,73 | 43.84 ± 3.58 | 1.01 ± 0.53<sup>a</sup> | 82 | 26.00 37.20<sup>a</sup> | 26.47 ± 19.15<sup>a</sup> | 11 |
|  | Medium | 9 | 6.15 ± 2.12 | 55.75 ± 6.92 | 1.29 ± 0.70<sup>a</sup> | 100 | 38.49 ± 55.87<sup>b</sup> | 57.62 ± 22.95<sup>ab</sup> | 12 |
|  | Large | 2 | 7.35 | 59.95 | 0.74<sup>ab</sup> | 100 | 15.00 ± 29.46<sup>ab</sup> | 47<sup>ab</sup> | 9 |
| Summer | Small | 20 | 3.43 ± 1.00 | 50.24 ± 5.44 | 2.50 ± 1.30<sup>b</sup> | 100 | 67.20 ± 34.21<sup>b</sup> | 100.06 ± 55.51<sup>a</sup> | 29 |
|  | Medium | 20 | 6.39 ± 1.41 | 62.52 ± 3.89 | 2.79 ± 0.65<sup>b</sup> | 100 | 66.40 ± 52.64<sup>b</sup> | 85.68 ± 27.97<sup>b</sup> | 29 |
|  | Large | 1 | 13.07 | 78.36 | 2.2<sup>ab</sup> | 100 | 49.03 ± 78.49<sup>b</sup> | 97<sup>ab</sup> | 9 |

**Figure 4.**
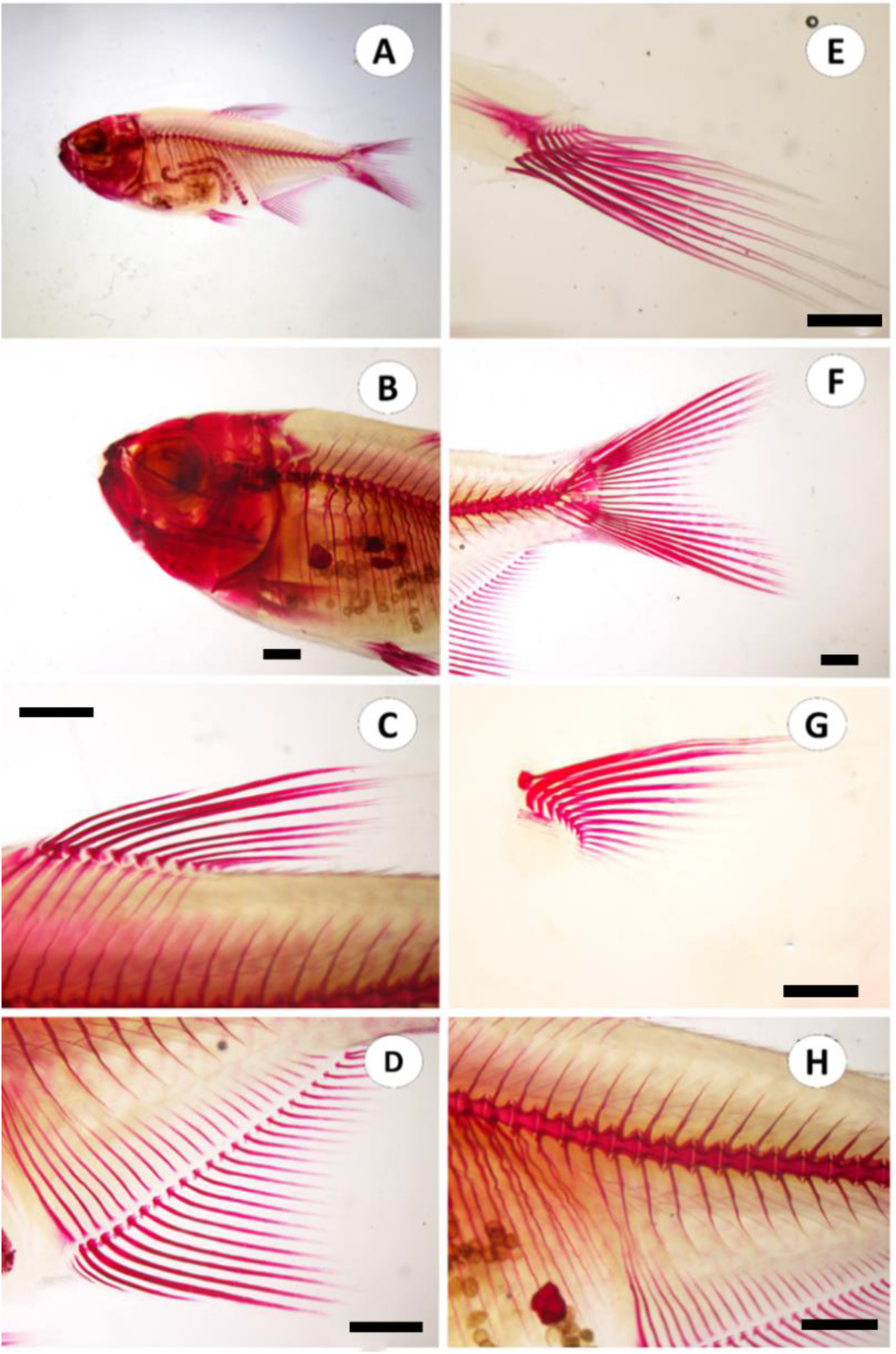
Fish diaphanized according to Potthoff (1984). Lateral view of *A. altiparanae* (A); Details of the cranial region (B); Dorsal fin (C); Anal fin (D); Pelvic fin (E); Caudal fin (F); Pectoral fin (G); (H) Dorsal spine. Scale bars: B-D, F, H: 1000 μm; E: 500 μm; G: 100 μm.

Only one female captured in summer presented spinelets, with a lower number and distribution when compared to males, and no female presented spinelets in winter (Table 3).

**Table III.** Meristic and somatic parameters of *Astyanax altiparanae* female in the Winter 2013 and Summer 2014.

| Season | Size classes | Number of animals (n) | Weight (g) | Pattern length (mm) | GSI (%) | Spinelets presence (%) | Spinelets length (mm) | Total of spinelets | Rays with spinelets |
| --- | --- | --- | --- | --- | --- | --- | --- | --- | --- |
| Winter | Small | 2 | 3.82 ± 0,58 | 46.66 ± 5.10 | 2.52 ± 1.38<sup>a</sup> | 0 | - | - | - |
|  | Medium | 16 | 7.27 ± 1.76 | 58.71 ± 6.02 | 3.20 ± 1.60<sup>a</sup> | 0 | - | - | - |
|  | Large | 23 | 17.67± 4.54 | 78.99± 7.18 | 3.20± 1.80<sup>a</sup> | 0 | - | - | - |
| Summer | Small | 5 | 2.2 ± 0.66 | 42.08 ± 3.93 | 1.60 ± 3.90<sup>a</sup> | 0 | - | - | - |
|  | Medium | 5 | 8.41 ± 0.96 | 65.54 ± 3.18 | 17.12 ± 4.67<sup>b</sup> | 0 | - | - | - |
|  | Large | 24 | 17.82 ± 3.28 | 83.34 ± 5.11 | 15.50 ± 7.5<sup>b</sup> | 4.17 | 117.89 ± 64.37 | 29 | 5 |

The GSI presented an increment from winter to summer due to the enter in the reproductive period. However, the females from small group showed no variation, suggesting that they had not reached their sexual maturation (Table 3).

### Experiment II

#### Long-term anesthesia for surgical procedures

The dosage of 0.1, 0.2, and 0.3% of 2-phenoxyethanol did not present efficacy, with just 0.78, 1.70, and 2.50 min of anesthesia, respectively. Those results were similar to that observed in the control (0.0%) with 0.90 min of anesthesia (P *< 0.0001*). On the other hand, with the dosage of 0.4% the time of anesthesia increased significantly (P = 0.0001) to 35.40 min, with one individual reaching 60 min. However, the dosage of 0.5% was sufficient for all fish remain in anesthesia for 60 min and also adequate for surgical procedures.

#### Gonadectomy

30 days after surgery, no differences were observed among control and gonadectomized fish for number (*P = 0.1122*; 70.20 ± 9.00 and 50.70 ± 8.50, respectively) (Fig. 5A) and rays with spinelets (*P = 0.7115;* 10.00 ± 1.00 and 9.20 ± 1.50, respectively) (Fig. 5B). However, a significant decrease (*P = 0.0478*) in the length of spinelets were observed for gonadectomized fish (90.82 ± 7.04 µm) in comparison to the control group (114.75 ± 8.26 µm) (Fig. 5C).

**Figure 5.**
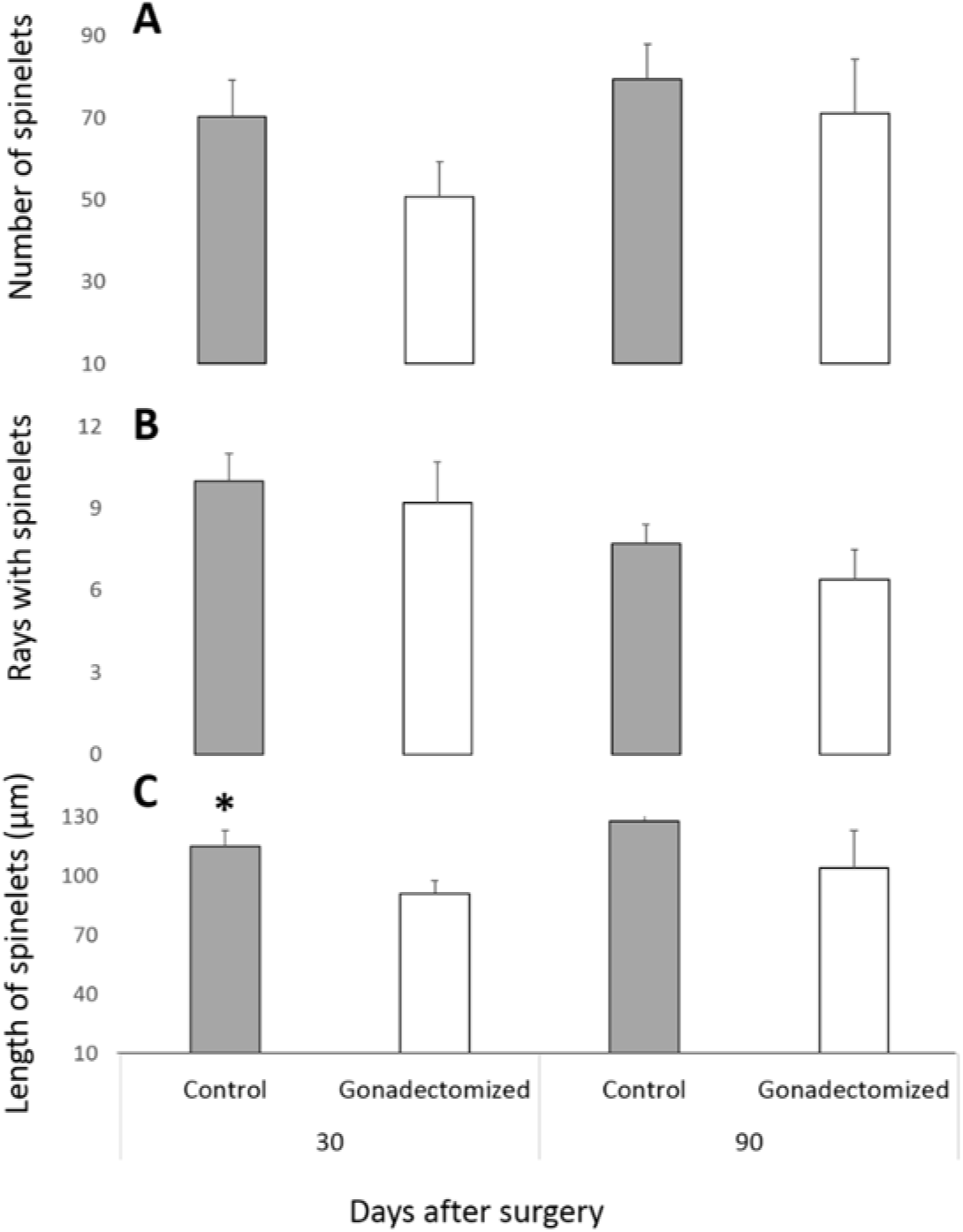
Comparison of the Number of spinelets (A), rays with spinelets; (B) and length of spinelets (µm); (C) between the control and fish with 30 and 90 days after surgery (gonadectomy) in *Astyanax altiparanae*.

90 days of surgery, no differences were observed among control and gonadectomized fish for number of spinelets (*P = 0.6138*; 79.30 ± 8.60 and 71.10 ± 13.10, respectively), rays with spinelets (*P = 0.2705*; 7.70 ± 0.70 and 6.40 ± 1.10, respectively) and length of spinelets *(P = 0.3554*; 127.70 ± 7.74 and 104.11 ± 18.89, respectively). One fish in gonadectomized group presented no anal fin spinelets.

## Discussion

The data presented show that the spinelets are not always a good parameter to differentiate gender in the yellowtail tetra *A. altiparanae*, as previously proposed (Siqueira-Silva et al., 2017). Moreover, their arise seems to be more linked to fish length and season of the year than to gonadal maturation phases, since spinelets were observed even in the Immature specimens collected during the summer. Some animals in Initial Maturation phases collected in winter had not. In media, the spinelets of the animals from the three categories (small, medium and large sizes) were bigger during the summer than those from the winter. However, it is important to highlight that, even in a moderate way, the gonads seems to influence the spinelets size as observed in thirty-day-gonadectomized-animals, in which spinelets were smaller in comparison to control fish.

The secondary sexual characteristics are very useful in fish management and breeding for many species. It can appear only during the breeding season, such the post occipital hum in *Cichla* species (Kullander & Ferreira, 2006), or being a permanently characteristic as in *Betta splendens* (Faria et al., 2006), in which males display a brightly colored body pattern. Generally, those characters are very important clues to ensure species reproductive success, which can be supplemented by courtship behavior and parental care. In other species, the difference is structural, such as in guppy (*Poecilia reticulata*), in which a copulatory organ, named gonopodium, develops from the male anal fin to be used in internal fertilization (Tian et al., 2015). The Atlantic salmon (*Salmo salar*) males use a hook developed on the lower jaw and enlarged teeth to perform kind of quiverings and yawning that seems to positively influence or initiate reproduction (Jarvi, 1990). Those characters, as the name suggest, are supposed to appear only during the reproductive season. The same was believed to the yellowtail tetra spinelets, which hypothetically were to develop only in males whose reproductive cycle have started. However, we found here that this character can be found even in Immature specimens during summer season. Thus, this characteristic cannot be used during the management to select potential male breeders during reproductive season. Moreover, since one spawning capable female also presented spinelets, the management to separate the genders in this species must be associated to other characteristics, such as the fusiform body in males and round body in females.

Similar data was described by Fujimoto and colleagues (2010), in the loach *Misgurnus anguillicaudatus.* In this species, a bony structure in the pectoral fin, named bony plate, was supposedly a phenotypical characteristic of sexually mature males only. However, such a structure was found in one wild fish showing an atypical like-ovary gonad. Moreover, as showed in our study, sterility did not influence the presence or absence of such a characteristic in the loach species. Muñoz et al. (2006) also found something similar in relation to the secondary sexual characteristic in the peacock bass *Cichla monoculus* when a single sampled female presented a post occipital hump, suggesting that energetic storage structure was always associated to sexually maturate males (Kelber, 1999). However, given the low number of occurrences in the above-mentioned species, this event might be an imbalance of the hormones in such specimens and must be faced as rare.

Finally, the reduction of spinelets size in this study might be related to a reduction of Estradiol (E2) synthesis, initially caused by gonadectomization of specimens, since this hormone is produced by Leydig Cells in the interstitial compartment of Teleosts. Estradiol hormone is stated to positively influence fish secondary sexual characteristics, as showed in guppy, whose animals incubated in E2 increased their gonopodium in relation to Control group.

In conclusion, this study showed that the yellowtail tetra (*A. altiparanae*) spinelets is a secondary sex characteristic influenced by sex, size, and season of the year more than by the gonadal development. However, as already showed in other species, such the loach (Fujimoto et al., 2010) and the pea spinelets arose even within gonadectomized fish, suggesting that such a structure is not an indicative of sterility in this species.

## Acknowledgments

We are grateful to the Sao Paulo Research Foundation (FAPESP) for the financial support of this research (2010/17429-1 and 2011/11664-1). We also acknowledge CEPTA/ICMBio for generously providing the facilities and experimental fish.

